# Tau-tubulin kinase 1 and amyloid-β peptide induce phosphorylation of collapsin response mediator protein-2 and enhance neurite degeneration in Alzheimer disease mouse models

**DOI:** 10.1101/855015

**Authors:** Seiko Ikezu, Kaitlin L. Ingraham Dixie, Lacin Koro, Takashi Watanabe, Kozo Kaibuchi, Tsuneya Ikezu

**Affiliations:** Departments of Pharmacology and Experimental Therapeutics, Boston University School of Medicine, Boston, MA, USA; Department of Cell Pharmacology, Graduate School of Medicine, Nagoya University School of Medicine, Aichi, Japan; Neurology, Boston University School of Medicine, Boston, MA, USA; Alzheimer’s Disease Center, Boston University School of Medicine, Boston, MA, USA; Center for Systems Neuroscience, Boston University School of Medicine, Boston, MA, USA

**Keywords:** Alzheimer’s disease, amyloid-β peptide, collapsin response mediator protein-2, entorhinal cortex, hippocampus, microtubule-associated protein tau, tau-tubulin kinase 1

## Abstract

The accumulation of phosphorylated tau protein (pTau) in the entorhinal cortex (EC) is the earliest tau pathology in Alzheimer’s disease (AD). Tau tubulin kinase-1 (TTBK1) is a neuron-specific tau kinase and expressed in the EC and hippocampal regions in both human and mouse brains. Here we report that collapsin response mediator protein-2 (CRMP2), a critical mediator of growth cone collapse, is a new downstream target of TTBK1 and is accumulated in the EC region of early stage AD brains. TTBK1 transgenic mice show severe axonal degeneration in the perforant path, which is exacerbated by crossing with Tg2576 mice expressing Swedish familial AD mutant of amyloid precursor protein (APP). TTBK1 mice show accumulation of phosphorylated CRMP2 (pCRMP2), in the EC at 10 months of age, whereas age-matched APP/TTBK1 bigenic mice show pCRMP2 accumulation in both the EC and hippocampal regions. Amyloid-β peptide (Aβ) and TTBK1 suppresses the kinetics of microtubule polymerization and TTBK1 reduces the neurite length of primary cultured neurons in Rho kinase-dependent manner *in vitro*. Silencing of TTBK1 or expression of dominant-negative Rho kinase demonstrates that Aβ induces CRMP2 phosphorylation at threonine 514 in a TTBK1-dependent manner, and TTBK1 enhances Aβ-induced CRMP2 phosphorylation in Rho kinase-dependent manner *in vitro*. Furthermore, TTBK1 expression induces pCRMP2 complex formation with pTau *in vitro*, which is enhanced upon Aβ stimulation *in vitro*. Finally, pCRMP2 forms a complex with pTau in the EC tissue of TTBK1 mice *in vivo*, which is exacerbated in both the EC and hippocampal tissues in APP/TTBK1 mice. These results suggest that TTBK1 and Aβ synergistically induce phosphorylation of CRMP2, which may be causative for the neurite degeneration and somal accumulation of pTau in the EC neurons, indicating critical involvement of TTBK1 and pCRMP2 in the early AD pathology.

## Introduction

Alzheimer’s disease (AD) is the most prevalent neurodegenerative disorder, characterized by the loss of synapses and neurons, leading to cognitive impairment and dementia. A growing body of evidence reveals that neurons affected in AD display “a dying-back pattern”, in which neurons start degenerating at the synapse or axon terminal then gradually die back toward cell-soma. Previous neuroimaging studies report significant axonal degeneration in the perforant path (PP) region in patients with mild cognitive impairment as determined by using magnetic resonance imaging and diffuse tensor imaging [1, 2]. The PP presents an important axonal tract that connects neurons in the entorhinal cortex (EC) to the dentate gyrus (DG) and other areas of the hippocampus [3]. This pathway is essential to many types of memory, including spatial memory [4], which is significantly impaired in AD patients. The axons of PP arise primarily from layer II neurons in the EC where somatic accumulation of hyperphosphorylated tau protein is detected in the early stage of AD. This dying-back axonal degeneration in the PP and accumulation of phosphorylated tau protein in the EC neuron are the critical initial events in AD pathophysiology in the context of the initiation and propagation of tau pathology. However, its mechanisms still remain elusive.

We have previously shown that tau-tubulin kinase1 (TTBK1), a brain-specific tau kinase, is specifically enriched in the PP of TTBK1 transgenic mice harboring BAC containing human *TTBK1* genome [5], followed by the EC, pyramidal layer of Cornu Ammonis (CA)1-3 fields and the granular layer of dentate gyrus in the hippocampal fields [6, 7]. A recent independent study has confirmed that TTBK1 mRNA is highly expressed in the human EC, subiculum, CA filed, and DG in both AD and non-demented cases [8]. TTBK1 directly phosphorylates the tau protein *in vitro*, and also enhances tau phosphorylation via unique activation of cyclin-dependent kinase 5 (Cdk5) upon dissociation of G-actin from Cdk5 *in vitro* and *in vivo* [6, 7, 5]. Further, TTBK1 protein expression is significantly up-regulated in frontal neocortical region of AD brain [7], and certain genetic variations of the TTBK1 gene are associated with late-onset AD in two cohorts of Chinese and Spanish populations [9, 10]. TTBK1 transgenic mice harboring the entire genomic sequence of human TTBK1 show accumulation of hyperphosphorylated tau in the cortex (including the EC) and hippocampal formation when crossed with P301L tau mice (JNPL3) [5]. Moreover, our recent study have shown that crossing P301L tau mice with TTBK1 transgenic mice accelerates axonal degeneration of spinal motoneurons, and silencing TTBK1 partially blocks lipopolysaccharide-stimulated microglia-induced neurite degeneration [11]. While multiple tau kinases including GSK3β, CDK5, or casein kinase 2 and tau itself have been reported to contribute to axonal degeneration in AD especially through axonal transport deficit [12], no kinase except TTBK1 is specifically expressed in the EC and PP where early AD pathology evolves. Thus, we hypothesize that TTBK1 may play a critical role in axonal degeneration of the PP through phosphorylation and aggregation of tau in AD.

In this study, we investigated the role of TTBK1 in AD pathology using an APP/ TTBK1 double transgenic mouse model. We specifically focused on the synergistic effect of TTBK1 and APP on phosphorylation of collapsing response mediator protein-2 (pCRMP2), an anterograde cargo transporter of tubulin dimer and a critical mediator of growth cone collapse that is also found in the neurofibrillary tangles [13], to elucidate the mechanism of axonal degeneration of the PP and accumulation of phosphorylated tau protein and pCRMP2 in the EC and hippocampal region in AD mouse models. AD patients’ brain samples were also tested to find the association between pathology measured on the Braak scale and TTBK1 and pCRMP2 expression in the EC and hippocampal region. Here we report that TTBK1 and CRMP2 may elucidate the relationship between Aβ deposition and EC-specific accumulation of phosphorylated tau protein, and neurite degeneration in early stage of AD.

## Materials and methods

### Transgenic animal models

All experimental procedures using animals were approved by the Institutional Animal Care and Use Committee at Boston University School of Medicine. Tg2576 (APP) mice expressing the Swedish mutation of human APP (isoform 695) were obtained from Drs. G. Carlson and K. Ashe and maintained in B6/129 background [14]. TTBK1 transgenic mice (Line 141) harboring human TTBK1 genomic fragment were maintained in B6/129 background as described previously [7, 5, 11]. Tg2576 mice were crossed with TTBK1 mice to generate APP, TTBK1, APP/TTBK1 and non-transgenic (non-Tg) littermates, and tested at 10-11 months of age.

### Immunofluorescence

Age-matched APP, TTBK1, APP/TTBK1 and non-Tg littermates (10-11 months of age) were deeply anesthetized with isoflurane inhalation and transcardially perfused with 4% paraformaldehyde in PBS (Sigma-Aldrich, St. Louis, MO). Brain tissues were cryoprotected by successive 24 h immersions in 15 and 30% sucrose in PBS, mounted in OCT compound. The tissue blocks were serially cryosectioned at 10 µm-thickness on coronal planes through regions of interest including the entorhinal cortex and hippocampal formations using a microtome cryostat (Microm HM525, Thermo Fisher Scientific), and mounted on Superfrost^TM^ Plus slides (Thermo Fisher Scientific). The tissue sections were subjected to immunofluorescence using pT514 CRMP2 rabbit polyclonal antibody (1:2000 dilution, generated by Kaibuchi laboratory), followed by Alexa Fluor 488 (Life technologies, Grand Island, NY) secondary antibody and Dapi (1:2000 dilution, Thermo Fisher Scientific). Immunostained images were captured using inverted fluorescence microscope attached to monochromatic CCD camera (TE-2000U, Nikon Instruments, Melville, NY) and fluorescence intensity were quantified with ImageJ software (NIH).

### Retrograde tracing by hippocampal injection of DiI

Paraformaldehyde-fixed mouse brain tissues were dissected in a coronal plane at -3.40mm (AP) posterior to bregma. The blocks including entorhinal cortex were post-fixed for at least 2 days in 4% paraformaldehyde in PBS. After post-fixation, 0.5 *μ*l of DiI (1,1’-Dioctadecyl-3,3,3’,3’-Tetramethylindocarbocyanine Perchlorate, 2.5 mg/mL in dimethyl sulfoxide, Thermo Fisher Scientific) were injected into the stratum moleculare of the dentate gyrus of the hippocampal filed using a motorized microinjector (Stoelting, Wood Dale, IL) and a 10-μL, gas-tight micro syringe (Hamilton, Reno, NV), which was set on the stereotaxic platform (David Kopf Instruments, Tujunga, CA). DiI injected tissues were stored in PBS and 0.2% sodium azide solution at 37°C for 4 weeks in the darkness. Tissue blocks were then embedded in 3% agarose gel and cut in the sagittal plane at 30 *μ*m-thickness by the vibratome (Leica, Buffalo Grove, IL), counter-stained with Dapi (1:2000 dilution, 20 min) and subjected to fluorescence microscopic imaging as described. DiI fluorescence intensity was quantified with ImageJ software (NIH).

### Plasmids and TTBK1 siRNA

DNA plasmids encoding HA-tagged human full length TTBK1 (HA-TTBK1) in pcDNA3.1+ vector was generated as previously described [6]. Myc-tagged human wild-type (WT) and mutant CRMP2 (S522A, T555A) [15] contained in a pCAGGS vector, and dominant negative RhoK [pEF-BOS-Myc-RB/PH (TT)] vector [16] were kindly provided by Dr Kaibuchi’s lab. Four putative TTBK1 siRNA sequences (clone 19; 5’-GCTCTTAAGGACGAAACCAACATGAGTGG-3’) were selected and subcloned into pVL-EGFP-Puro vectors. Upstream of the green fluorescence protein (GFP) coding sequences, siRNA sequences were inserted into pSuper siRNA vectors at the Hind III restriction site.

### Tissue and Cell culture

Human neuroblastoma cell line SH-SY5Y that stably expressed wild type Tau (2N4R, 1-411; kindly provided by Roger M. Nitsch) [17, 18] or human embryonic kidney 293 cell line (HEK293) cells were maintained in a 5% CO2 humid atmosphere at 37°C in Dulbecco’s modified eagle medium/Ham’s F12 nutrient mixture (DMEM/F12) with 10% fetal bovine serum, 10 U/ml penicillin, and 10 µg/ml streptomycin (all from Life technologies) as described [6]. Primary cultured murine cortical neurons were prepared from E16 embryonic CD-1 mouse brains and plated on poly-D-lysine-coated coverslips in 24-well plates (Thermo Fisher Scientific Inc., Waltham, MA) as described [11].

### DNA plasmid transfection and amyloid-β treatment

HEK 293 cells (2 × 10^6^ cells / 6 well plate) were transfected with 2 μg of plasmids (CRMP2 wild type, CRMP2 S522A mutant, or CRMP2 T555A mutant) or 20 pmol of siRNA with 10μl Lipofectamine 2000 (Thermo Fisher Scientific). Twenty-four h after the transfection, cells were incubated with 0, 3, or 10 μM of freshly prepared human synthetic amyloid-β peptide 1-42 (Aβ42, Life technologies) for 24 h as described (Kiyota et al., 2011). Primary murine cortical neurons (1 × 10^5^ cells/24 well plate) were cultured as previously described [11] and transfected with 0.5 μg each of plasmids [TTBK1, GFP and/or dominant negative RhoK] with 4 μL of Lipofectamine 2000 (Life technologies).

### Analysis of neurite length and branches

Three days after the transfection, murine cortical neuronal cells were fixed with 4% paraformaldehyde in PBS solution, and GFP signal was captured by monochromatic CCD camera. NeuronJ plug-in was used on ImageJ (NIH, Bethesda, MD) to measure the length of axons and number of dendrites emanating from the cell soma [19].

### Immunoblotting

Whole cell lysates were prepared in RIPA lysis buffer containing 1% NP-40, 0.5% sodium deoxycholate and 0.1% SDS in the presence of 1 mM PMSF, 1 mM NaF and 1 mM Na_3_VO_4_ after centrifugation at 20,817 *× g* for 30 min at 4°C. Twenty μg protein/sample were subjected to 4-15 % SDS-PAGE (Bio-Rad laboratories, Hercules, CA) and immunoblotted for anti-phosho-Thr514 CRMP2 (pT514-CRMP2, rabbit polyclonal, 1:1000) [20], anti-total CRMP2 mouse monoclonal (C4G mAb, 1:500, IBL. Inc, Minneapolis, MN) [21], and anti-TTBK1 mouse monoclonal (20 μg/ml, clone F287–1.1–1E9) [7].

### End-binding protein 3 (EB3)-GFP Live Confocal Microscopy

SH-SY5Y cells plated on 35 mm MatTek dishes (MatTek Corp, Ashland, MA) at 2 × 10^5^ cells / dish and transfected with EB3-GFP plasmid (0.5 μg) was co-transfected with TTBK1 or pcDNA3.1^+^ plasmids (0.5 μg) as described [22, 23]. The live imaging of comet movement of EB3-GFP homodimer was captured 24h later by the following setting: 100*×* oil objective (NA=1.45), 488 nm excitation (100-300 ms exposure/frame, single plane), 0.53 fps for 40 frames, using the Nikon TE-2000U inverted fluorescent microscope (Nikon Instruments, Melville, NY) output port and cooled charge-coupled device camera (Coolsnap HQ, Roper Scientific, Duluth, GA) as described [24, 23].

### Immunoprecipitation assay

Recombinant Myc-tagged human CRMP2 was transiently expressed in SH-SY5Y cells stably expressing P301L human tau (2N4R) [18], and treated with 10 μM of freshly prepared amyloid-β peptide 1-42 (A*β*42, Life technologies) for 24 h. Forty eight h after the DNA plasmid transfection, cell lysates were prepared using RIPA buffer and incubated over night at 4°C with anti-Myc rabbit polyclonal (1:150 dilution, Covance Inc, Dedham, MA) or anti-phosho-T514 CRMP2 rabbit polyclonal antibody (1:100 dilution, generated by Kaibuchi Laboratory). The mixture was then incubated with 20 μL of 50% protein G plus-agarose (Sigma-Aldrich, St. Louis, MO) for 4 h at 4°C with gentle rocking. After three washes in the ice-cold RIPA buffer, the immunoprecipitates were separated by 4-10% SDS-PAGE (Bio-Rad) and immunoblotted for phosphorylated tau (AT8 mouse monoclonal, 1:500 dilution, Thermo Fisher Scientific), unphosphorylated tau (Tau-1 mouse monoclonal, 1:200 dilution, Millipore, Temecula, California), total tau (Tau46 mouse monoclonal, 1:500 dilution, Abcam, Cambridge, MA), or total CRMP2 (clone C4G, mouse monoclonal, 1:500 dilution, IBL America, Minneapolis, MN).

For *in vivo* tau and CRMP2 binding assay, the cortical and hippocampal tissues were dissected from mouse brains and homogenized in ice-cold RIPA buffer containing protease and phosphatase inhibitor cocktails (Roche), followed by centrifugation at 20,817 *× g* for 30 min at 4°C. The protein concentration of the protein extract supernatant was determined using a BCA assay kit (Thermo Fisher Scientific). pCRMP2 was immunoprecipitated from 2 mg of protein extracts by anti-phosho-T514 CRMP2 rabbit polyclonal antibody (1:100 dilution) and protein A/G sepharose (Santa Cruz Biotech, Dallas, TX), and the immunoprecipitants were blotted for phosphorylated tau (AT8) and total tau (Tau46).

### Human brain specimens

The entorhinal cortex and hippocampus brain tissue sample were obtained from the Boston University AD Center Brain Bank and also the Harvard Brain Tissue Resource Center. Post mortem interval was < 24 h, if it was possible, with the brain samples from Boston University Brain Bank. The criteria for AD were based on the presence of amyloid-β neuritic plaques and neurofibrillary tangles according to the NIA–Reagan criteria for intermediate and high likelihood AD and the recent NIA Alzheimer Association’s guidelines [25, 26]. The NIA–Reagan criteria take into account both the Braak staging for neurofibrillary tangles [27, 28] and the overall density of neuritic plaques based on CERAD criteria [29]. The hippocampal and entorhinal cortex tissues were obtained from the Braak stage I-IV patients. The demographic information of all of the patients from Boston University AD Center Brain Bank used for this study are described in Table 1. Brain tissues were paraffin embedded, then cut at 8 μm by the microtome by Boston University Pathology Laboratory Service Core. The paraffin sections were mounted on Superfrost^TM^ Plus slides (Thermo Fisher Scientific).

**Table 1.** ********.

| Patient ID | Age (YR) | Braak<br>Stage | Sex | TTBK1 |  | pCRMP2 |  |
| --- | --- | --- | --- | --- | --- | --- | --- |
|  |  |  |  | EC II | CA1 | EC II | CA1 |
| 1 | 89 | I | M | - | - | + | +++ |
| 2 | 87 | I | F | + | + | - | - |
| 3 | 92 | I | F | + | + | - | - |
| 4 | 90 | II | M | ++ | ++ | ++ | ++ |
| 5 | 84 | II | M | - | - | ++ | +++ |
| 6 | 96 | II | F | + | + | - | - |
| 7 | 93 | III | M | ++ | + | ++ | +++ |
| 8 | 93 | III | F | + | ++ | + | +++ |
| 9 | 92 | III | F | + | + | +++ | + |
| 10 | 92 | IV | M | + | + | +++ | +++ |
| 11 | 96 | IV | F | + | + | ++ | +++ |
| 12 | 81 | IV | M | +++ | +++ | +++ | +++ |

### Immunohistochemistry (human brain sections)

Deparaffinized sections were incubated with proteinase K (20 *μ*g/ml) in Tirs-EDTA-CaCl_2_ buffer at 37 °C in humidified chamber and room temperature for 10 min each, rinsed in tris-buffered saline (TBS) with 0.025 % Triton X-100 and permeabilized with 0.1% Triton-X 100 in TBS for 10 min. After further washing with TBS, sections were incubated with blocking buffer (10% horse serum with 1% bovine serum albumin in TBS) for 2 h at room temperature, then incubated with following primary antibodies at 4 °C for 48 h: TTBK1 mouse monoclonal (1:20, clone 1E9 generated by Ikezu laboratory) [7] and 3F4 anti-pCRMP2 mouse monoclonal (1: 100, ILB America), which detects pT509/pS518/pS522 [21]. Endogenous peroxidase activity was inhibited with 0.3% hydrogen peroxide (Sigma-Aldrich) incubation. Sections were then incubated with secondary anti-mouse antibody (Immpress-HRP anti-mouse IgG (peroxidase polymer detection kit, Vector Laboratories, Burlingame, CA) for 30 min. The 3,3′-Diaminobenzidine (DAB; Vector Laboratories) staining was used as a chromogen, and hematoxylin was used for counterstaining. Immunostained images were captured using a Nikon Eclipse E600 microscope and a color charge-coupled device camera (Nikon Instruments, Melville, NY). The 3F4 pCRMP2^+^ cells were quantified using 10 sections/donor in the EC II region, and ranked as – (< average 0.5 cell/field), + (average 0.5 – 1 cell/field), ++ (average 1– 2 cell/field), +++ (average >2 cell/field).

DAB intensity of TTBK1 in EC and CA1 regions was measured according to the previous methods [30]. Briefly, captured images were saved as 24-bit RGB TIFF format and converted to CMYK TIFF format by adobe Photoshop CS5 software. Intensity in the Y channel of individual cells was measured as DAB^+^ immunoreactive signals by image J software. Signal/Noise (S/N) ratio of at least 50 cells were averaged and ranked as – (≤ 1 S/N), + (1 < S/N ≤2), ++ (2 < S/N ≤3), +++ (3 < S/N ≤4), and ++++ (4 < S/N). Representative transformation from original RGB images to Y channel images in CMYK format was shown in Supplementary Figure 1.

## Results

### TTBK1 and phosphorylated CRMP2 are highly expressed in the EC and hippocampal region in AD brains

In order to investigate the involvement of TTBK1 in AD pathology at the early stage, we assessed whether the expression of TTBK1 in the EC and hippocampal region is correlated with Braak stage of AD using human brain specimens (Table 1). Immunohistochemistry using TTBK1 monoclonal antibody (clone IE9) revealed that TTBK1 is expressed highly in the cell soma and dendrites in the EC and hippocampal neurons including the DG, subiculum and CA1-3 field (Fig. 1A). Intensity measurements of TTBK1 immunoreactivity in the pyramidal cell soma of layer II EC neurons and CA1 showed consistent expression of TTBK1 from Braak I to IV stages (Table 1). In addition, TTBK1^+^ neurons were partially co-localized with AT8^+^specific to pS202/pS205 phosphorylated tau (pTau) in the frontal cortex in AD patients’ brains (Fig. 1B), indicative of expression of TTBK1 in tau-bearing neurons in these regions.

**Figure 1.**
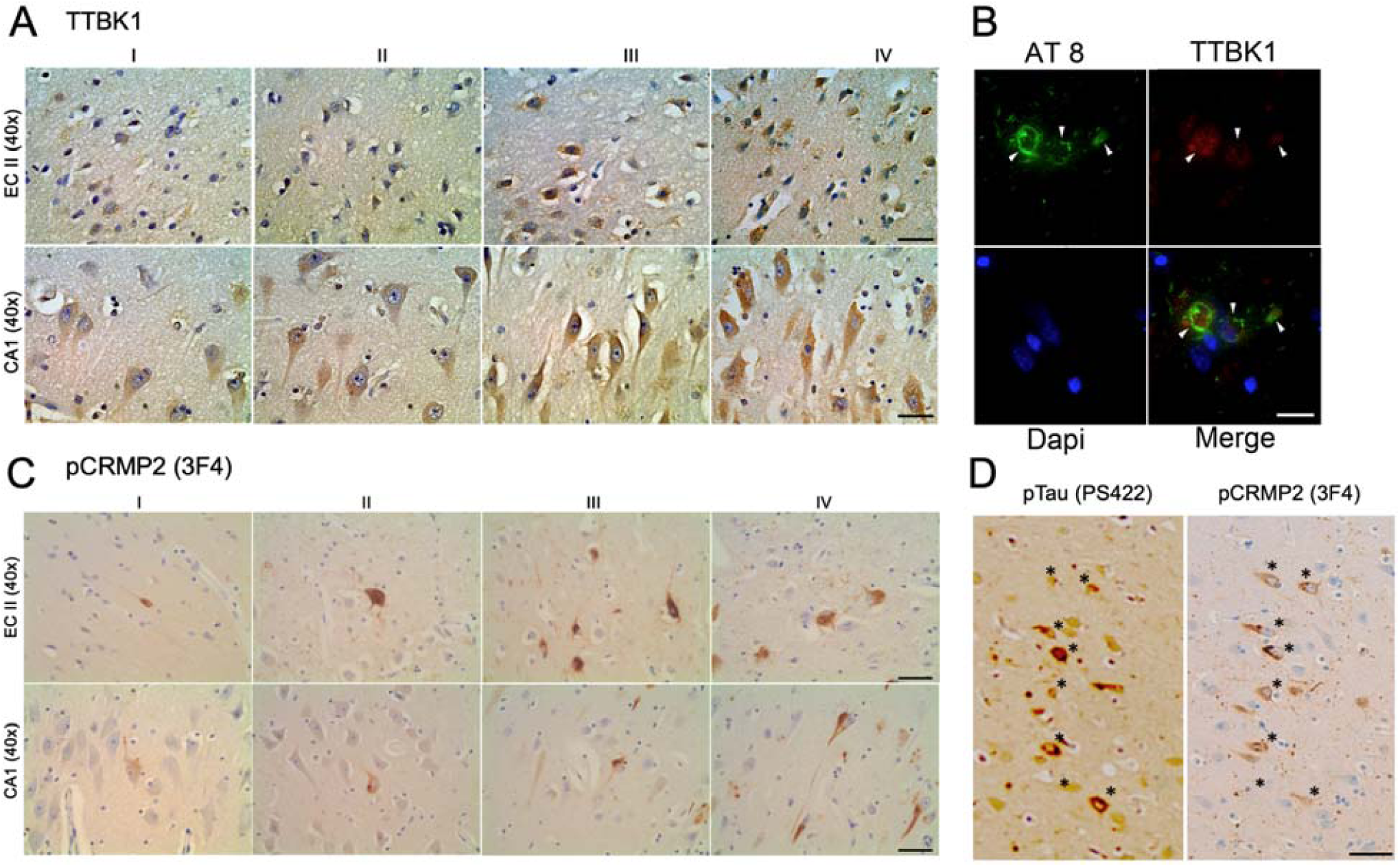
Localization of TTBK1 and pCRMP2 in the pyramidal neurons of the EC and CA1-3 hippocampal regions in early AD brains. The human brain specimens are listed in Table 1. **A,** Deparaffinized human brain sections (8 *μ*m in thickness) containing the EC layer II and CA1 hippocampal regions from patients in Braak I-IV were DAB immunostained for TTBK1 and counterstained with hematoxilyn. Top panel: EC regions, bottom panel: CA1 region. Scale bar = 20 *μ*m. **B,** Immunofluorescence of Braak II EC brain sections with AT8 (pSer202/205 tau, green), TTBK1 (red) and Dapi (blue). Scale bar = 10 *μ*m. **C,** DAB immunostaining with 3F4 (pCRMP2) in the EC layer II and CA1 hippocampal regions from patients in Braak I-IV. Scale bar = 20 *μ*m. **D,** Co-localization of 3F4 (pCRMP2) and pSer422 Tau in the AD brain sections (indicated by *). Scale bar = 40 *μ*m.

Immunohistochemistry against 3F4 detecting phosphorylated CRMP2 (pCRMP2) at pT509/pS518/pS522 showed pCRMP2^+^ neurons as early as Braak I in the EC, subiculum and hippocampal CA1-3 fields and we found that the number of pCRMP2^+^ neurons were correlated with Braak stages (Fig. 1C and Table 1). In addition, we observed co-localization of pCRMP2 and pS422 pTau in the CA1 field as determined by immunostaining of the adjacent sections of human brain tissue with 3F4 (for pCRMP2) and PS422 (for pTau, Figure 1D). Pervious study reported that colocalization of TTBK1 with PS422 in pre-tangle neurons determined by immunohistochemistry with AD and non-symptomatic control brain samples[8], indicating a synergistic effect with TTBK1 and pCRMP2 in early tau pathology.

These data demonstrate that both TTBK1^+^ and pCRMP^+^ are expressed in the EC and hippocampal neurons in AD patients’ brains from the early Braak stage, and pTau^+^ neurons show co-localization with TTBK1 or pCRMP2, suggesting the potential contribution of TTBK1 in pCRMP2 and pTau accumulation.

### Synergistic enhancement of pCRMP2 accumulation by TTBK1 and APP expression in the mouse brain

To assess whether increased expression of TTBK1 and Aβ accumulation directly induces pCRMP2 accumulation *in vivo*, we crossed Tg2576 mice expressing KM670/671NL Swedish mutation of human APP_695_ (APP mice) [14] and TTBK1 mice expressing human *TTBK1* from bacterial artificial chromosome [7] to generate TTBK1, APP, APP/TTBK1, and non-Tg littermates. These mice were tested at 10-11 months of age for immunofluorescence against pT514-CRMP2 in the EC and hippocampal regions. pT514-CRMP2 was detected in the neurons of layer II/III of the EC of APP, TTBK1 and APP/TTBK1 mice but not in non-Tg mice (Fig. 2A, green). APP/TTBK1 mice show the most pronounced accumulation of pCRMP2 in the EC region, followed by TTBK1 and APP mice (Fig. 2B). These data support the previous *in vivo* demonstration of the age-dependent accumulation of pCRMP2 in the cortical region of Tg2576 [31]. Further, APP/TTBK1 mice also showed accumulation of pCRMP2 in the granular cell layer of the dentate gyrus (DG), suggesting the synergistic effect of APP and TTBK1 expression in this region (Fig. 2C-D, green). CA field did not show any pCRMP2 accumulation with these 3 mouse models. Taken together, we observed expression of pCRMP2 in the EC in aged TTBK1 and AD mouse models, which was exacerbated in APP/TTBK1 mice, whereas pCRMP2 accumulation in the DG was only observed in APP/TTBK1 mice.

**Figure 2.**
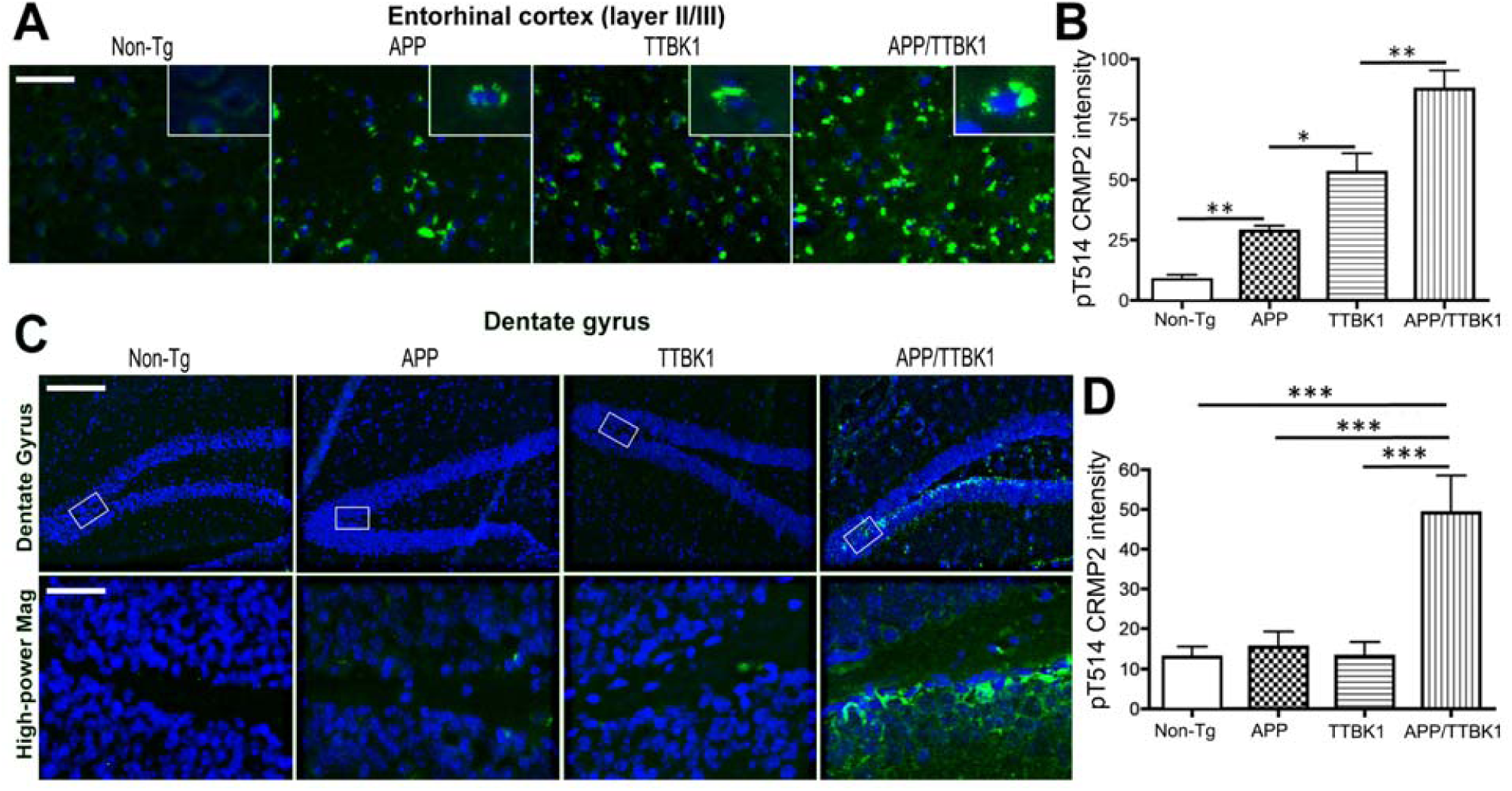
Transgenic expression of APP and TTBK1 synergistically enhance accumulation of pCRMP2 in the entorhinal cortex and dentate gyrus. APP (Tg2576), TTBK1, APP/TTBK1 and age-matched non-Tg mice at 10-11 months of age were subjected to immunofluorescence against pCRMP2 (T514, green) and Dapi (blue). **A-B,** Layer II/III of the EC (A) was quantified for pCRMP2 intensity (B). Insets in A represent pCRMP2 accumulation in the neuronal cell soma. **C-D,** pCRMP2 staining was also observed in the subgranular zone and mossy fiber regions of the dentate gyrus of APP/TTBK1 mice but not in the other groups. Lower panel represents 20x original magnification of the subgranular zone. *, **, and *** denotes p < 0.05, 0.01, or 0.001 as determined by one-way ANOVA and Tukey *post hoc* (N = 8 per group). Scale bars represent 50 *μ*m (A), 200 *μ*m (C, upper panel), and 30 *μ*m (C, lower panel).

### Axonal degeneration in the perforant path (PP) of TTBK1 and APP transgenic mice

We have previously reported the specific accumulation of TTBK1 immunoreactivity in the PP of TTBK1 mice and significant accumulation of phosphorylated neurofilament heavy chain in the EC layer II/III, which is indicative of axonal degeneration [7]. Accumulation of pT514-CRMP2 is also indicative of the growth cone collapse *in vitro* [15, 32, 20]. In order to investigate the involvement of TTBK1 and Aβ in the axonal degeneration of the PP, we tested the retrograde transport of DiI from the DG to the EC region through the PP. Retrograde tracer DiI was injected into the outer molecular layer of the DG of the hippocampal region of TTBK1, APP, APP/TTBK1 and non-tg littermates, and the accumulation of DiI in the PP and EC regions was examined. Interestingly, we observed significant reduction of DiI signals in the PP and layer II/III region of the EC in TTBK1 mice and, to the lesser extent, APP mice compared to age-matched non-tg littermates (Fig. 3A-B). The reduction of DiI signals was exacerbated in APP/TTBK1 mice compared to either TTBK1 or APP mice. Thus, we found that the intensity of DiI signal in the EC region was the highest in the non-tg group, followed by APP, TTBK1 and APP/TTBK1 mice (Fig. 3C-D). Taken together, these data demonstrate that TTBK1 and APP expression individually induces axonal degeneration of EC neurons in the PP, and that co-expression of TTBK1 and APP further enhances the axonal degeneration. Importantly, axonal degeneration in the PP are positively correlated with pCRMP2 accumulation in the EC as previously tested with those mouse models(Fig. 2B), suggesting the orchestrated effect by TTBK1, Aβ and CRMP2 phosphorylation leading to compromised axonal integrity.

**Figure 3.**
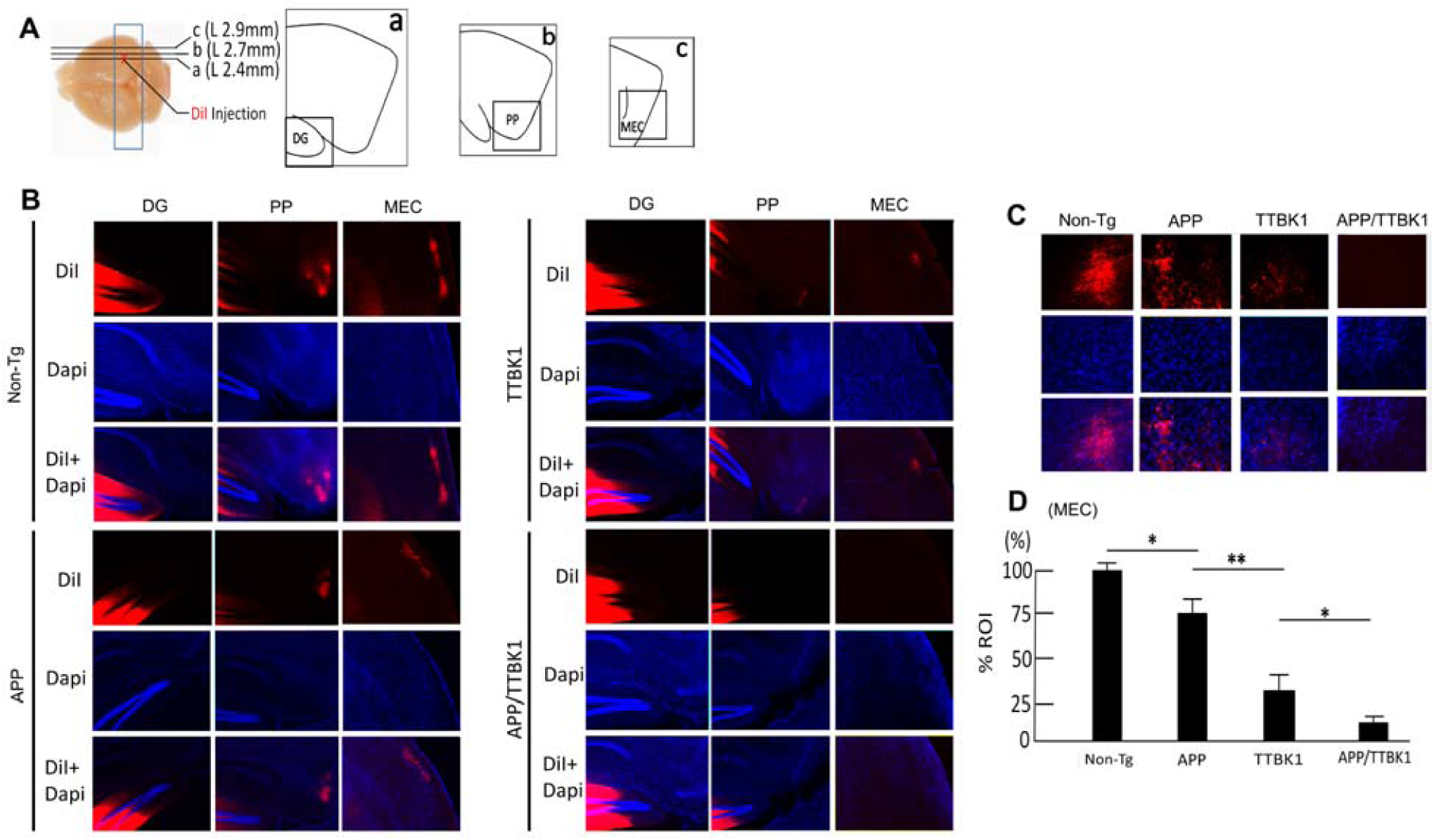
Transgenic expression of APP and TTBK1 induces axonal degeneration of EC neurons. **A,** Stereotaxic DiI injection into stratum moleculea of dentate gyrus of a fixed and vibratome sectioned mouse brain . Transgenic littermates were sacrificed at 10 months of age and fixed brain tissues were injected with 0.5 μl of DiI into the stratum moleculae of dentate gyrus (DG). DG (a), PP (b) and MEC (c) regions were illustrated from the sections proximal to ML +2.4 (a), +2.7 (b) and +2.9 (c). **B**, DiI-labeled axon tracts (red) counterstained with Dapi (blue) in the DG, PP, and MEC in Non-Tg, APP, TTBK1, and APP/TTBK1 mice. **C,** Retrograde tracing of DiI in the layer II/III region of EC of transgenic littermates. **D,** DiI intensity measurements in the region of interest (ROI) of the layer II/III region of MEC. * denotes p < 0.05 as determined by one-way ANOVA and Tukey *post hoc* (N = 6 per group).

### Aβ and TTBK1 reduce microtubule polymerization kinetics

The collapsin molecules, such as semaphorin or Eph family, mediate growth cone collapse through phosphorylation of CRMP2 [15]. CRMP2 is a cargo molecule to carry α/β-tubulin hetero-dimers to the tip of microtubules via its binding to kinesin [33]. The collapsin-mediated phosphorylation of CRMP2 leads to the dissociation between itself and the tubulin dimers and kinesin, which triggers growth cone collapse and neurite retraction [34]. Given that Aβ and TTBK1 may not only induce phosphorylation of tau but also potentially CRMP2, a key molecule to maintain the neurite integrity, we assessed if Aβ and TTBK1 has effect on the dynamics of microtubule assembly, which may contribute to the neurite destabilization observed in TTBK1, APP, and APP/TTBK1 mice. We employed confocal microscopic real-time live imaging of microtubule polymerization by monitoring transiently expressed microtubule plus-end-binding (EB) 3-EGFP in SH-SY5Y human neuronal cells. EB3-GFP forms a homodimer at the plus-end tips of polymerizing microtubules, which enabled us to analyze the velocity of microtubule polymerization (Fig. 4A-B) [24]. Treatment of EB3-GFP-expresssing SH-SY5Y cells with freshly prepared Aβ42 oligomer or co-expression of TTBK1 significantly reduced the EB3-GFP comet velocity by 28 or 13% as compared to the mock-transfected control, respectively (Fig. 4C). These data demonstrate that Aβ42 oligomer and TTBK1 can suppress the neurite elongation via suppression of the velocity of microtubule polymerization, which may lead to impaired neurite stability.

**Figure 4.**
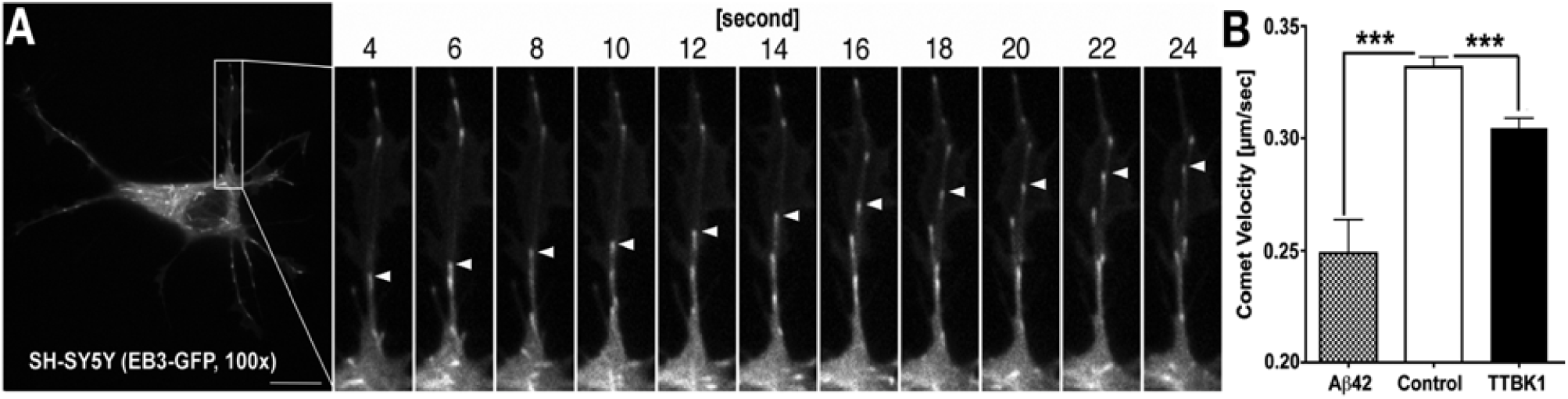
Time-lapse image of microtubule polymerization. **A,** SH-SY5Y cells were transfected with EB3-GFP fusion vector, and laser scanning live confocal images were taken 24 h after DNA transfection at 0.53 fps for 40 frames with Em/Ex 488/520 nm using 100*×* Oil objective (NA=1.45). Arrowheads indicate one of the EB3-GFP comets polymerizing outward from the microtubules. Scale bar: 5 µm. **B,** Velocities of EB3 comets were quantified after co-transfection of EB3-EGFP (0.2 µg) with/out HA-TTBK1 (0.4 µg), followed by treatment with 10 µM freshly dissolved A*β*42 at 1 h prior to the imaging. *** denotes p < 0.001 by one-way ANOVA with Turkey *post hoc* (N = 10 cells per group).

### TTBK1 expression and Aβ stimulation induces CRMP2 phosphorylation at T514 *in vitro*

We have previously shown that TTBK1 activates Cdk5 via dissociation of Cdk5 from G-actin and recruitment of p35/p25 to Cdk5 [7, 35]. Other evidence has shown that Cdk5 directly phosphorylates CRMP2 at Ser522, which is the priming step of the phosphorylation of CRMP2 at threonine 514 (T514) by GSK3β and this phosphorylation of CRMP2 at T514 is critical for the induction of neurite retraction [20]. We thus examined if TTBK1 expression facilitates phosphorylation of CRMP2 at T514 via Cdk5 activation *in vitro*. We found that transient expression of TTBK1 in human embryonic kidney 293 (HEK293) cells induced significant phosphorylation of CRMP2 at T514 (Fig. 5A, lane 1 and 2). This was abolished by Ser 522 to Ala mutation (S522A, lane 3 and 4) at the Cdk5 phosphorylation site, and partly abolished by Thr 555 to Ala mutation (T555A, lane 5 and 6) at the Rho kinase (RhoK) phosphorylation site [36, 32, 20]. These data suggest that TTBK1-induced T514 CRMP2 phosphorylation is dependent on both S522 phosphorylation by Cdk5 and T555 phosphorylation by RhoK.

**Figure 5.**
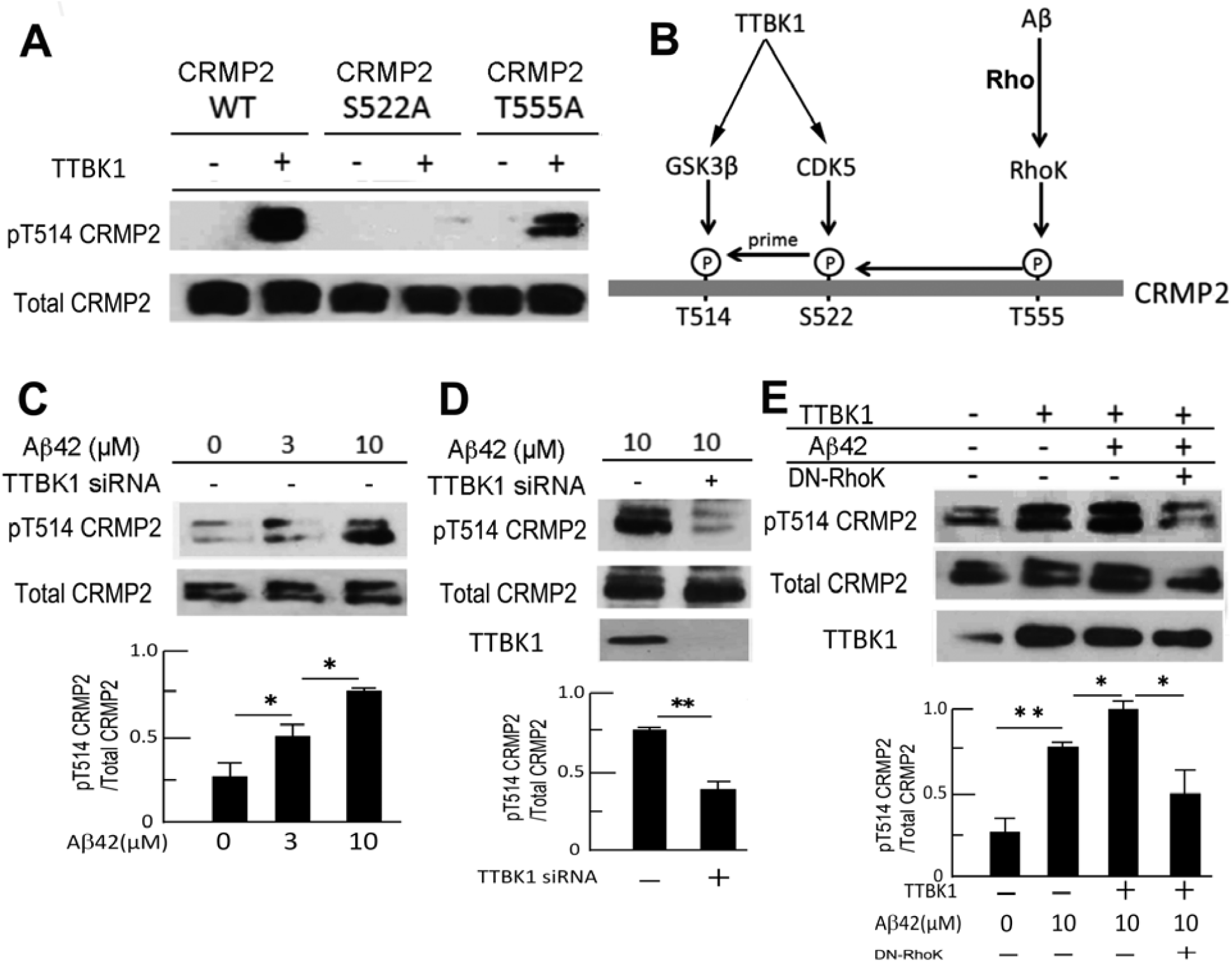
Aβ and TTBK1 induces CRMP2 phosphorylation *in vitro*. A. HEK293 cells were transfected with CRMP2 expression plasmids [wild type (WT), S522A mutant, and T555A mutant] with or without human wild type TTBK1 plasmid, followed by immunoblotting of total and phosphor-CRMP2 at pT514. **B,** Schematic diagram of site-specific CRMP2 phosphorylation by TTBK1 and Aβ42. **C-D**, SH-SY5Y cells were transfected with scramble or TTBK1 siRNA, followed by stimulation with different doses of freshly prepared Aβ42 (0, 3, and 10 µM) for 24 h. Cell lysates were subjected to immunoblotting of endogenous pCRMP2 at T514, and endogenous TTBK1. **E,** SH-SY5Y cells were transfected with a combination of TTBK1 and DN-Rho plasmids (0.5 µg each), followed by stimulation with 10 µM freshly prepared Aβ42 for 24 h. Results were representative of 3 independent experiments. * and ** denote p < 0.05, or 0.01 as determined by one-way ANOVA and Tukey *post hoc* (N = 3 per group).

In addition to evidence of the relationship between TTBK1 and pCRMP2, previous reports suggest that Aβ induces pCRMP2 in a RhoK-dependent manner [37]. Using human neuronal SH-SY5Y cells, we found that Aβ42 stimulation phosphorylates endogenous CRMP2 at T514 in a dose-dependent manner (Fig. 5C). Interestingly, we found that Aβ42-induced phosphorylation of CRMP2 at T514 is dependent on TTBK1, since the siRNA-mediated silencing of endogenous TTBK1 significantly reduces the pT514-CRMP2 level by 80% as compared to controls (Fig. 5D). Moreover, we observed synergistic enhancement of pT514-CRMP2 levels by Aβ treatment and TTBK1 expression, which was significantly reduced by the co-expression of dominant-negative RhoK (DN-RhoK) (Fig. 5E). These data demonstrate that Aβ42-induced pT514-CRMP2 is dependent on TTBK1, and that synergistic phosphorylation of pCRMP2 by Aβ42 and TTBK1 is dependent on RhoK (Fig. 5B).

### TTBK1 induces neurite degeneration in a Rho-dependent manner

Recent studies have shown that phosphorylation of CRMP2, an anterograde cargo transporter of tubulin dimer, is involved in the neurite degeneration pathology in multiple neurodegenerative diseases [38][39]. Indeed, several axon guidance molecules mediate their signaling through phosphorylation of CRMP2 [40]. Thus, based on our findings, we hypothesized that TTBK1 and Aβ suppress neurite elongation by CRMP2 phosphorylation, which is dependent on Rho-mediated RhoK activation. To test this hypothesis, primary cultured mouse cortical neurons were transiently transfected with TTBK1 and green fluorescent protein (GFP). Expression of TTBK1 dramatically shortened axon length compared to controls (Fig. 6A-B). Since Aβ and TTBK1-induced T514 CRMP2 phosphorylation is dependent on its T555 phosphorylation by RhoK, and Rho activates RhoK to phosphorylate CRMP2 at T514 [15], we examined if the Rho/RhoK pathway is involved in TTBK1-mediated neurite degeneration by co-transfection of dominant negative Rho (DN-Rho). The reduced axon length was restored by co-transfected DN-Rho, while single transfection of DN-Rho had no effect as compared to the GFP control group (Fig. 6B). Similarly, TTBK1 expression reduced the number of primary branches of dendrites, which is partially restored by the co-transfection of DN-Rho (Fig. 6C). These data indicate that TTBK1 expression suppresses neuronal neurite extension and dendritic branching partially through the Rho/RhoK pathway.

**Figure 6.**
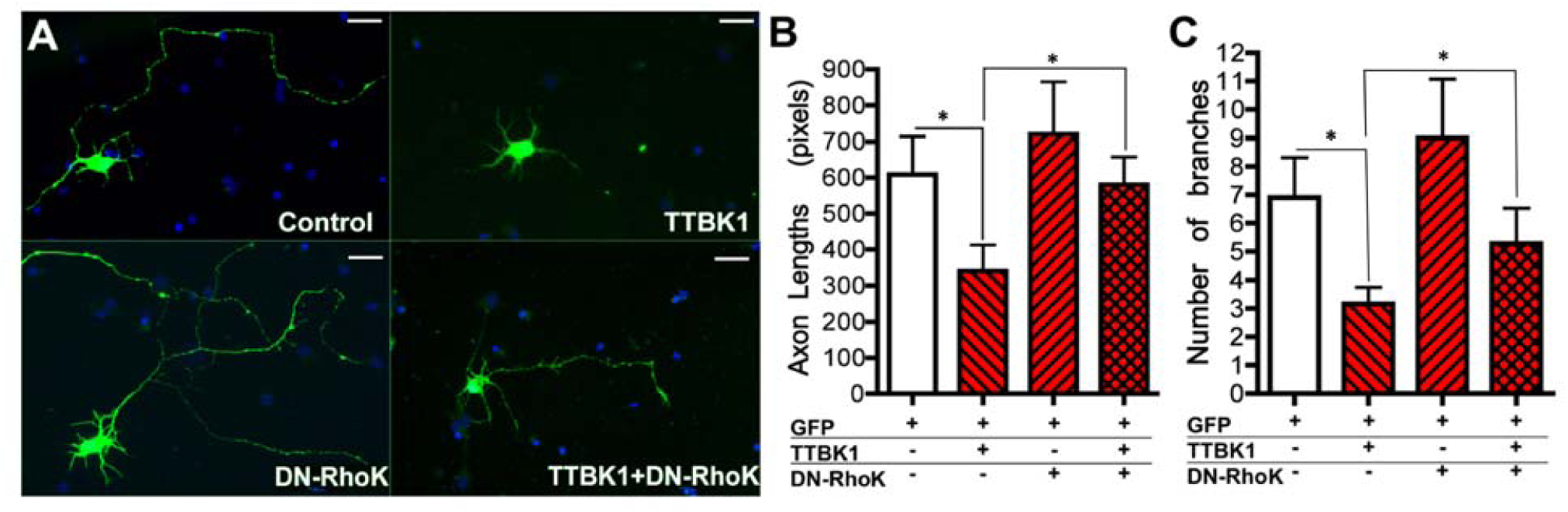
TTBK1 suppresses axonal elongation in Rho-dependent manner. **A,** Primary cultured mouse cortical neurons (green) were cultured on the PDL-coated coverslip in 24-well tissue culture plate, transfected with DNA plasmids expressing GFP, TTBK1, and/or DN-Rho (0.5 µg) at 5 div and fixed at 8 div. Blue represents Dapi counterstain. **B-C,** The axonal length (B) and the number of branches from primary axons (C) in GFP^+^ neurons were determined. Results were representative of 3 independent experiments. * denotes p < 0.05 as determined by one-way ANOVA and Tukey *post hoc* (N = 10 cells per group). Scale bars represent 100 *μ*m.

### TTBK1 induces complex formation of pCRMP2 with pTau

Since TTBK1 phosphorylates CRMP2 and tau proteins, and pCRMP2 is a component of neurofibrillary tangles in the AD brain [21], we hypothesized that TTBK1 expression or Aβ treatment directly induces physical association of CRMP2 and tau protein in their phosphorylation-dependent manner. To assess this hypothesis, Myc-tagged CRMP2 were expressed with or without TTBK1 in SH-SY5Y cells stably expressing human full-length wild type tau protein [17], followed by treatment with freshly prepared Aβ42. The cell lysate samples were immunoprecipitated for CRMP2 by anti-Myc antibody, and immunoblotted for pTau (AT8), total tau (Tau46), and non-phosphorylated tau (Tau-1). TTBK1 expression significantly induces CRMP2 binding to tau, which was enhanced by Aβ42 treatment (Fig. 7A-B, top panels), while Aβ42 treatment itself, conversely had no effect on the binding (Fig. 7A-B, top panels). The physical association of CRMP2 to tau appears to be dependent on tau phosphorylation, since no Tau-1 (specific to non-phosphorylated tau) immunoreactivity was observed in CRMP2-immunoprecipitated samples (Fig. 7A, top panel). A similar finding was also observed in the samples immunoprecipitated with PT514-CRMP2 antibody and immunoblotted for AT8, Tau46, and Tau-1 (Fig. 7A middle panel, 7B bottom panel). The immunoblot of the 5% of the input for the immunoprecipitation shows equal amount of total tau (T46) and CRMP2 expression, induction of AT8^+^ pTau by TTBK1 expression and its enhancement by A*β*42 stimulation (Fig, 7A, bottom panel). Tau-1 immunoreactivity, representing non-phosphorylated tau, is reduced by TTBK1 expression and A*β*42 treatment. These data demonstrate that TTBK1 expression is sufficient for CRMP2/tau complex formation in their phosphorylation dependent manner, and Aβ42 stimulation can augment the complex formation.

**Figure 7.**
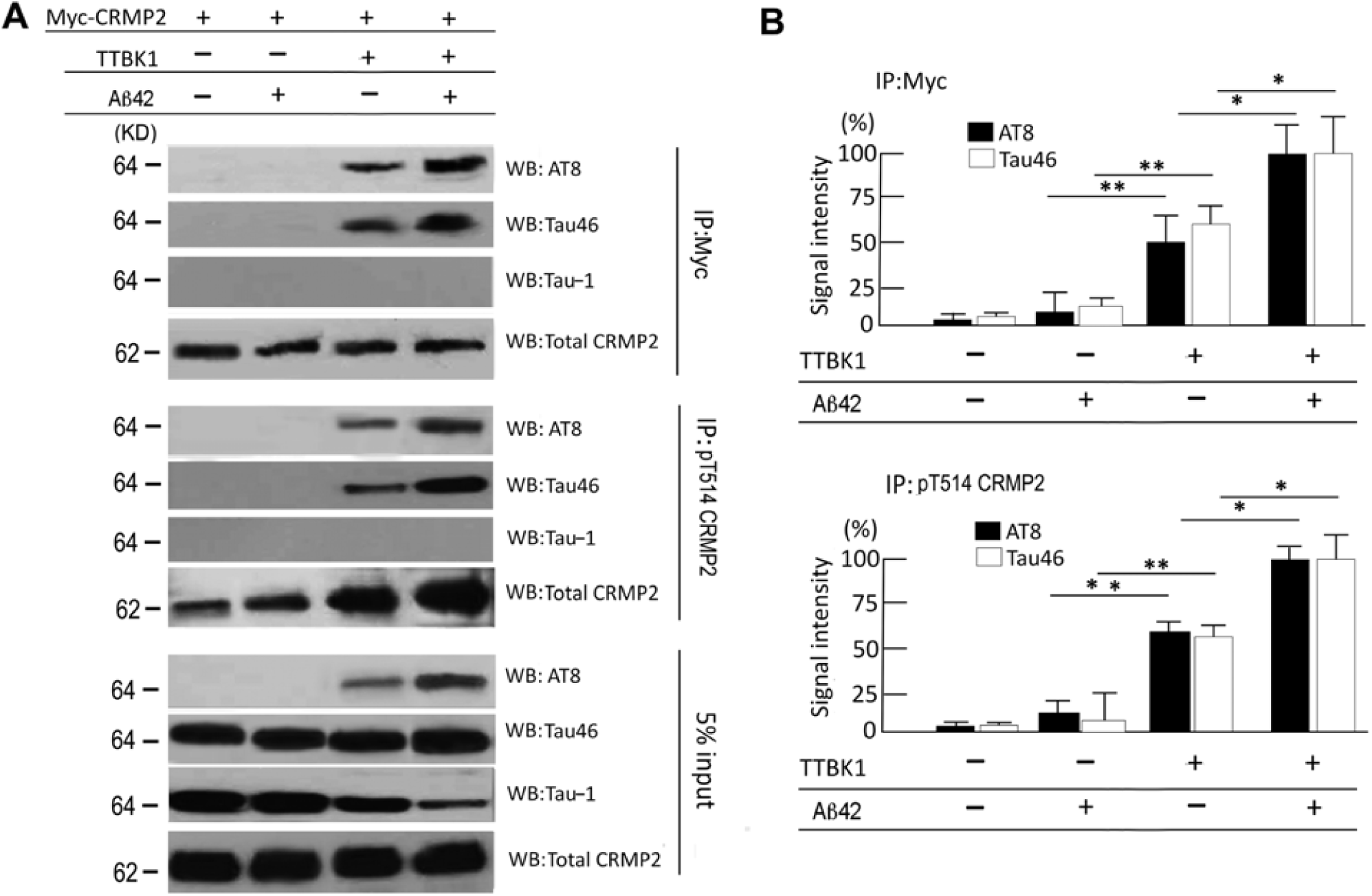
TTBK1 induces physical association of CRMP2 and tau *in vitro*. **A,** SH-SY5Y cells were transfected with plasmids expressing human wild type tau (2N4R), Myc-tagged WT CRMP2 with or without human wild type TTBK1 (0.5 µg each). Cells were treated with 10 µM freshly prepared Aβ42. The cell lysates were immunoprecipitated with anti-Myc or pT514-CRMP2 antibody, followed by immunoblotting of the immunoprecipitated samples with AT8 (anti-pS202/pS205 pTau), Tau-1 (unphosphorylated tau), Tau46 (total tau) and total CRMP2 (C4G). 5% of input lysate was also blotted for tau (Tau46) and β-actin for loading normalization. **B,** Quantification of AT8 or Tau46-immureactive bands after normalization with co-precipitated pCRMP2 levels (top panel) or pT514 CRMP2level (bottom panel) (N = 3 per group). Results were representative of 3 independent experiments. * and ** denote p < 0.05, or 0.01 as determined by one-way ANOVA and Tukey *post hoc* (N = 3 per group).

### pCRMP2 binds to pTau in the EC of TTBK1 and APP/TTBK1 mouse brain

To determine whether TTBK1 induces pTau and pCRMP2 complex formation *in vivo*, we assessed accumulation of pTau in pCRMP2-positive neurons in the specific brain regions in TTBK1 and APP mouse models. Our attempt to detect pTau in the EC or hippocampal region by AT8 or PHF-1 staining in these mice was unsuccessful at this age. This suggests that the endogenous tau expression is insufficient for detecting the pTau/CRMP2 complex formation by immunofluorescence. Therefore, brain lysates from the EC and DG with TTBK1 and APP mouse models were immunoprecipitated for T514-phosphorylated CRMP2 and immunoblotted for tau (Fig. 8A-B). pCRMP2 forms a complex with pTau in the EC regions of TTBK1 and APP/TTBK1 mice, while this complex formation in the hippocampal region was only detected in APP/TTBK1 mice (Fig. 8A). The immunoreactivity of co-precipitated tau (AT8 or Tau46) was strongest in APP/TTBK1, followed by TTBK1, APP mice and non-Tg littermates in the EC (Fig. 8B), whereas it was only significant in the hippocampus with APP/TTBK1 mice (Fig. 8B). Taken together, these data demonstrate that TTBK1 but not APP transgene expression alone induce the complex formation of pTau and pCRMP2 in the EC, whereas co-expression of APP and TTBK1 transgene enhance the complex formation of pTau and pCRMP2 in both the EC and hippocampal regions.

**Figure 8.**
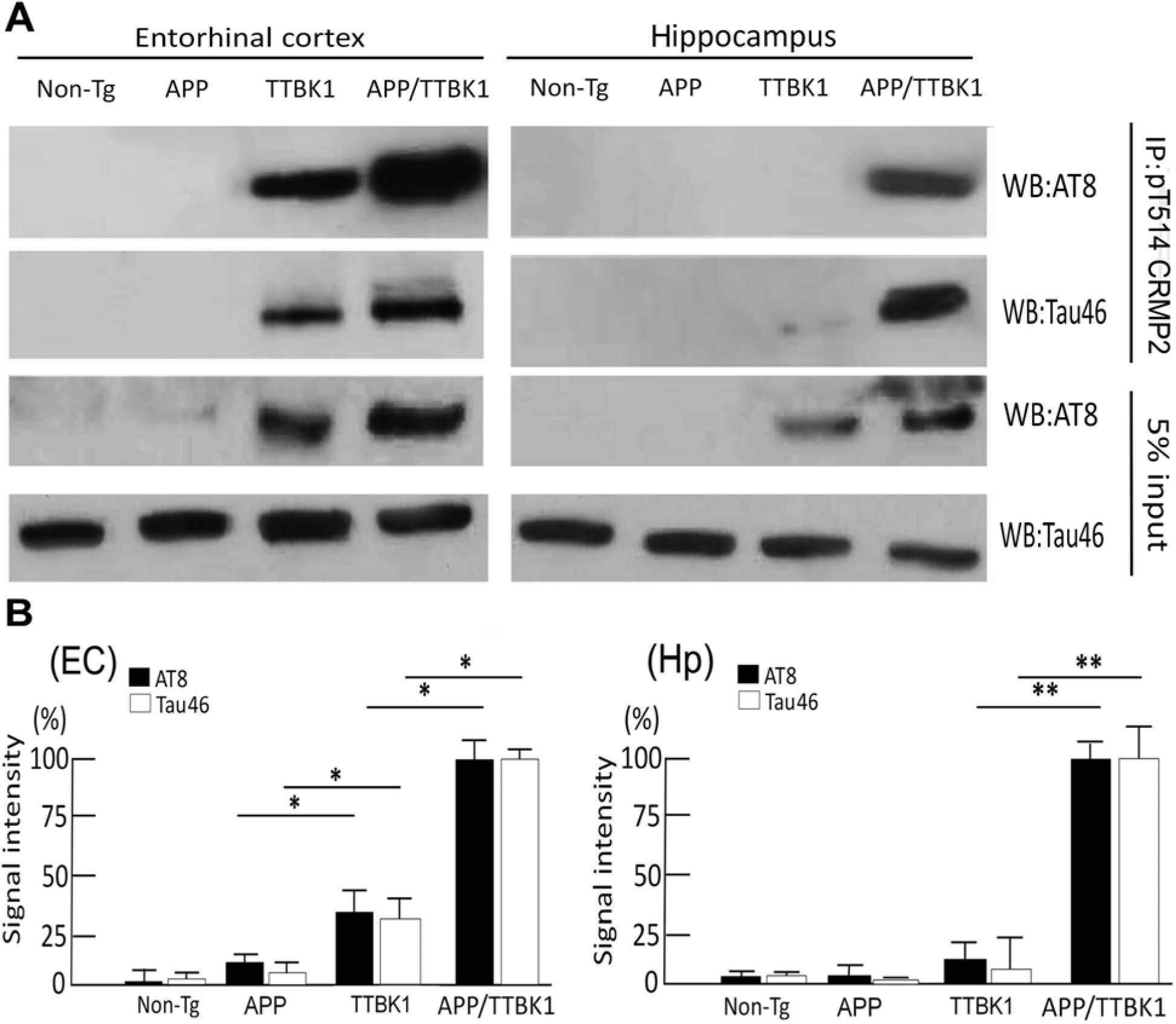
Transgenic expression of APP and TTBK1 induces physical association of CRMP2 and tau *in vivo*. **A,** EC and hippocampus lysates were immunoprecipitated with anti-pT514-CRMP2 antibody, followed by immunoblotting of the immunoprecipitated samples with AT8 (anti-pS202/pS205 pTau) or Tau46 (total tau). Five % of input lysate was also blotted for AT8 and total tau (Tau46). **B,** Quantification of AT8 or Tau46-immureactive bands after normalization with co-precipitated pT514 CRMP2level from EC (left panel) or hippocampal tissues (Hp, right panel). B. Signal intensity was quantified. * and ** denote p < 0.05, or 0.01 as determined by one-way ANOVA and Tukey post hoc (N = 6 per sample).

## Discussion

In this study, we have demonstrated that 1) the expression of TTBK1 and accumulation of pCRMP2 is observed in the EC, subiculum, and CA1-3 pyramidal neurons of AD brains, and the number of pCRMP2 is correlated with Braak scale for AD tau pathology, 2) TTBK1, APP and TTBK1/APP mice show somal accumulation of T514 phosphorylated CRMP2 and axonal degeneration in EC neurons, 3) TTBK1 and Aβ reduces the velocity of microtubule extension, 4) TTBK1 expression suppresses neurite elongation and branching, 5) Aβ and TTBK1 induces phosphorylation of T514 CRMP2, which is dependent on its S522 and T555 phosphorylation and 6) TTBK1 induces phosphorylation of CRMP2 and tau, and their complex formation *in vitro* and *in vivo and* Aβ enhances its effect.

TTBK1 mice show significant axonal degeneration in the PP as determined by accumulation of phosphorylated neurofilament heavy chain in the EC region [7]. In this study, we directly assessed the axonal degeneration in the PP via DiI-tracing from the DG to EC, which was evident in TTBK1 mice and enhanced in APP/TTBK1 mice. Accumulation of pT514-CRMP2 in the EC neurons was associated with axonal degeneration observed in APP, TTBK1 and APP/ TTBK1 mouse models, therefore we hypothesized that axonal degeneration facilitated by TTBK1 are mediated by pCRMP2. CRMP2 plays a physiological role on neurite elongation by transporting *α*/β-tubulin heterodimer to the axonal terminal via kinesin-1-dependent anterograde transport [33, 41]. CRMP2 is also an essential molecule for regulating the axon elongation by axon guidance molecules (such as Semapholin 3A) and Aβ [40, 31], and its collapsing function is regulated by its site-specific phosphorylation at S555, S522 and finally T514 by RhoK, Cdk5 and GSK3β, respectively [15, 32, 20]. Therefore, the lack of tubulin heterodimer supply to the plus-end of the microtuble terminal by pCRMP2 may initiate microtubule depolymerization, leading to the neurite degeneration (Fig.9).

**Figure 9.**
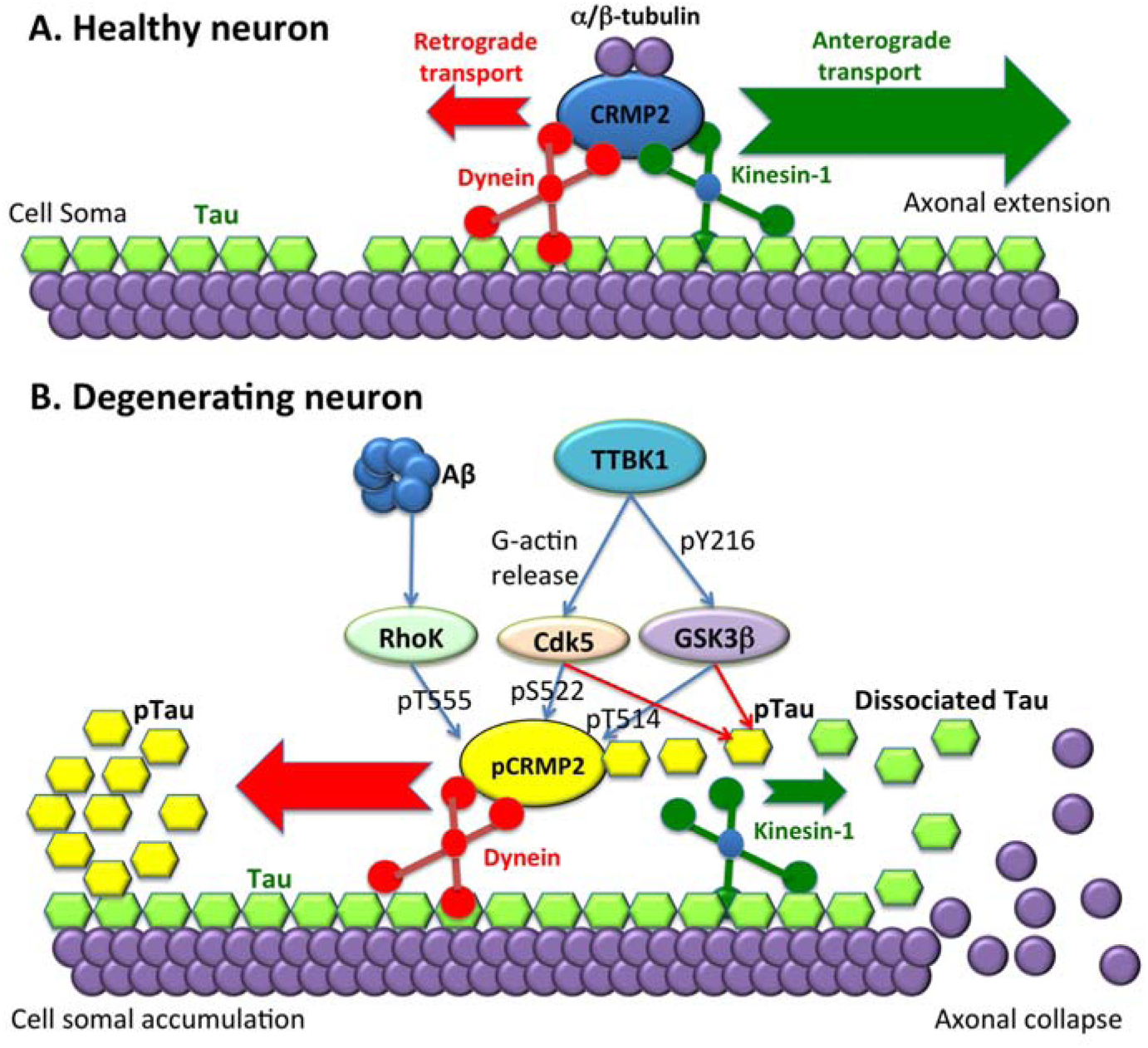
Schematic diagram of Aβ and TTBK1-induced phosphorylation of CRMP2 and tau, neurite degeneration and their accumulation in neuronal cell soma. **A,** In healthy neurons, kinesin-1-mediated anterograde transport is dominant (green arrow) over dynein-mediated retrograde transport (red arrow) for transporting α/β-tubulin dimer for axonal extension. **B,** Upon stimulation with Aβ or activation of protein kinases by TTBK1, CRMP2 is phosphorylated by Cdk5, GSK3β and RhoK (pCRMP2, yellow), and dissociates from kinesin-1, leading to more retrograde-dominant transport (red arrow) over their anterograde transport (green arrow). The loss of supply of α/β-tubulin by CRMP2 leads to depolymerization of neurite terminal, which dissociates tau (green). Dissociated tau is phosphorylated by tau kinases (pTau, yellow), which binds to pCRMP2 and concentrates in the cell soma via retrograde transport. The pCRMP2/pTau complex was found in EC neurons in TTBK1 mice. Accumulation of Aβ may accelerate this process and promote pCRMP2 accumulation in not only EC but also DG neurons in APP/TTBK1 mice.

Interestingly, recent studies suggest involvement of pCRMP2 in axonal degeneration pathology in multiple sclerosis both in an animal model and human brain samples [38]. pCRMP2^+^ axonal staining was found in the plaque core and peripheral white matter with acute and chronic-active multiple sclerosis patients. A similar result was also found in the spinal cord injury of rat model and the experimental autoimmune encephalomyelitis mouse model. In this study, we identified a unique signaling pathway where TTBK1 induces CRMP2 phosphorylation at pT514 that is dependent on its phosphorylation at S522 and T555. Moreover, we also found that Aβ induces pT514-CRMP2 in a TTBK1 and RhoK-dependent manner. We showed that both Aβ and TTBK1 suppress microtubule polymerization as determined by EB3-GFP live imaging and the neurite degeneration in TTBK1-transfected mouse cortical neurons in a Rho-dependent manner. Taken together, these data suggest that Aβ induces Rho activation and pT514-CRMP2 leading to neurite degeneration in a TTBK1-dependent manner.

A previous study and ours showed that pCRMP2 binds to pTau [42]. Further, phosphorylation of CRMP2 was identified in the neurofibrillary tangles isolated from AD brain [21]. We have shown for the first time that pCRMP2 is co-localized in phospho-Ser 422 positive pyramidal CA1 neurons with Braak Stage 1 human brain tissue. Recent work has shown that TTBK1 is co-localized with pS422^+^ pTau in neuronal cell soma containing pre-tangles but not in neuropil threads or neurofibrillary tangles [8]. Since TTBK1 can directly phosphorylate tau at S422, this data suggest its role in pS422^+^ pTau accumulation in pre-tangle^+^ neurons [6, 43]. In this study, we confirmed that TTBK1 is co-localized with AT8^+^ pTau in neurons, supporting the previous finding of our data showing that TTBK1/P301L tau bigenic mice exhibited enhanced AT8^+^ tau oligomer formation compared to P301L tau mice [5]. Moreover, p-CRMP2 and p-tau complex formation was induced by TTBK1 and exacerbated by Aβ *in vitro* and *in vivo*. Therefore, these findings suggest that together with TTBK1, pCRMP2 may be involved in the early phase of tau phosphorylation and pre-tangle formation in AD pathology.

In conclusion, our study has integrated the biological function of CRMP2, region-specific expression and molecular function of TTBK1 and Aβ accumulation as novel pathogenic mechanism of neurite degenerations and tau accumulation in the affected brain regions of AD, most importantly in the EC layer II. Since individuals in Braak stage I or II are prodromal AD and mostly cognitively normal, early intervention of neurite degeneration by TTBK1-specific inhibitors may potentially serve as a preventive therapy for AD. Unlike RhoK, GSK3β, or Cdk5, TTBK1 is specifically expressed in neurons and thus it is an attractive therapeutic target with less off-target potential.

## Acknowledgements

We would like to thank Drs. H. Asai and other members of Laboratory of Molecular NeuroTherapeutics or supporting this research, Dr. Y. Ihara (Doshisha University) for his suggestion of TTBK1 CRMP2 connection, Dr. Y. Goshima (Yokohama City University) for CRMP2 plasmids, Dr. K. Ashe (University of Minnesota) for Tg2676 mice, Dr. S. Popov (University of Illinois at Chicago) for EB1- and EB3-GFP plasmids and Ms. S. Walsh (University of Nebraska Medical Center) for assistance in live confocal microscopic imaging and analysis.

## Authors’ Declaration

The authors express no conflict of interest in this manuscript

The authors declare the data and material availability in this manuscript

---

**Supplementary Figure S1.**
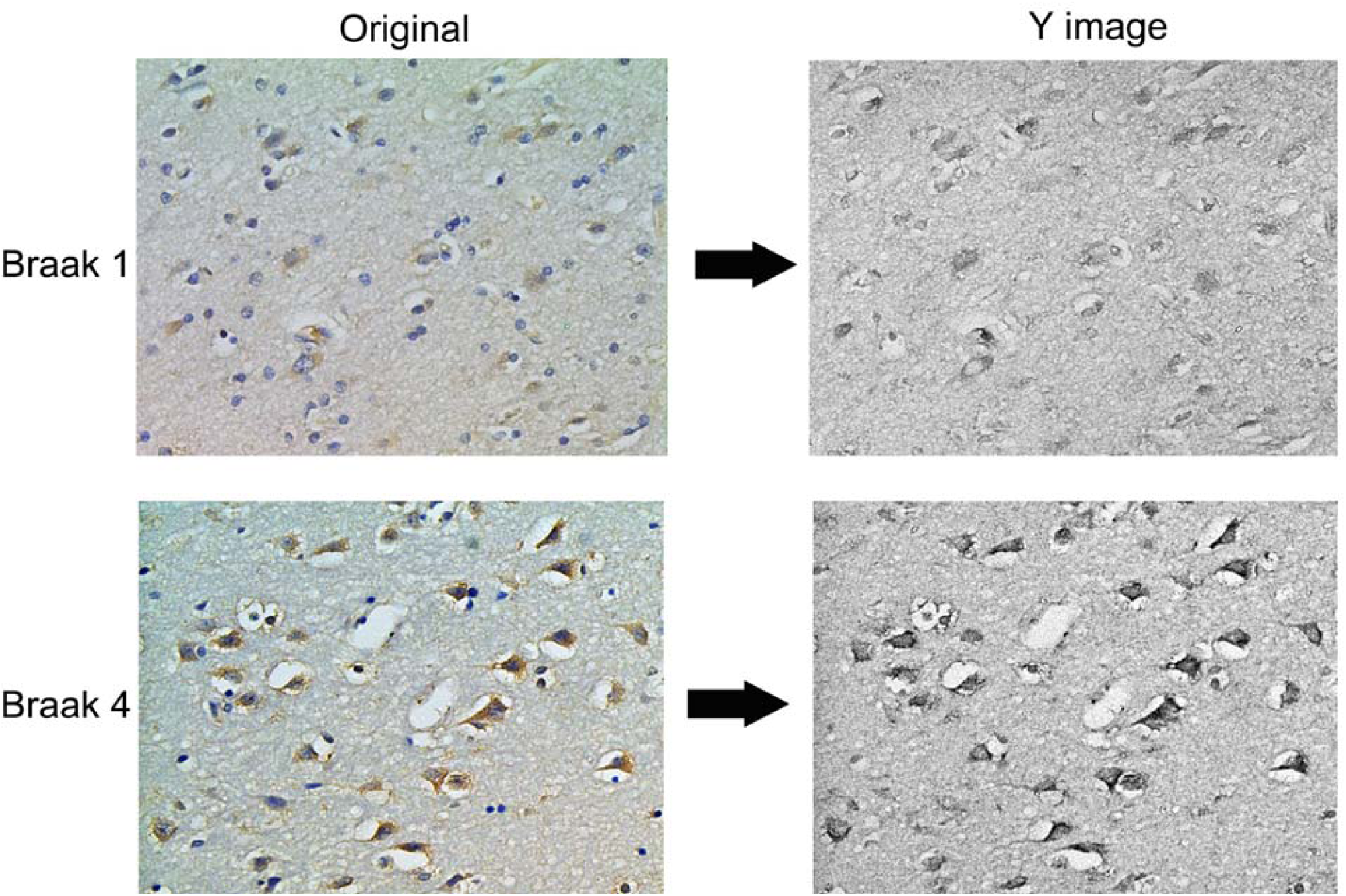
Representative conversion of CMYK format and extraction of Y channel (right) for DAB intensity measurement.

